# Similarity of brain activity patterns during learning and subsequent resting state predicts memory consolidation

**DOI:** 10.1101/2020.09.04.283002

**Authors:** Z. Zavecz, K. Janacsek, P. Simor, M.X. Cohen, D. Nemeth

**Affiliations:** Doctoral School of Psychology, ELTE Eötvös Loránd University, Izabella utca 46., H-1064 Budapest, Hungary; Institute of Psychology, ELTE Eötvös Loránd University, Izabella utca 46., H-1064 Budapest, Hungary; Brain, Memory and Language Research Group, Institute of Cognitive Neuroscience and Psychology, Research Centre for Natural Sciences, Magyar tudósok körútja 2., H–1117 Budapest, Hungary; Centre of Thinking and Learning, Institute for Lifecourse Development, School of Human Sciences, University of Greenwich, Old Royal Naval College, Park Row, 150 Dreadnought, SE10 9LS London, United Kingdom; Institute of Behavioural Sciences, Semmelweis University, Üllői út 26, H-1085 Budapest, Hungary; UR2NF, Neuropsychology and Functional Neuroimaging Research Unit at CRCN - Center for Research in Cognition and Neurosciences and UNI - ULB Neurosciences Institute, Université Libre de Bruxelles (ULB), 50 ave. F.-D. Roosevelt CP191 B-1050 Brussels, Belgium; Donders Centre for Medical Neuroscience, Radboud University Medical Center, Geert Grooteplein Zuid 10, 6525 Nijmegen, the Netherlands; Lyon Neuroscience Research Center (CRNL), Claude Bernard University Lyon 1, Centre Hospitalier Le Vinatier - Bâtiment 462 - Neurocampus 95 boulevard Pinel 69675 Bron, France

**Keywords:** memory consolidation, EEG, functional connectivity, similarity, resting state, reactivation, reinstatement, procedural learning, statistical learning, sequence learning

## Abstract

Long-term memory depends on memory consolidation that seems to rely on learning-induced changes in the brain activity. Here, we introduced a novel approach analyzing continuous EEG data to study learning-induced changes as well as trait-like characteristics in brain activity underlying consolidation. Thirty-one healthy young adults performed a learning task and their performance was retested after a short (~1h) delay, that enabled us to investigate the consolidation of serial-order and probability information simultaneously. EEG was recorded during a pre- and post-learning rest period and during learning. To investigate the brain activity associated with consolidation performance, we quantified similarities in EEG functional connectivity of learning and pre-learning rest (baseline similarity) as well as learning and post-learning rest (post-learning similarity). While comparable patterns of these two could indicate trait-like similarities, changes in similarity from baseline to post-learning could indicate learning-induced changes, possibly spontaneous reactivation. Individuals with higher learning-induced changes in alpha frequency connectivity (8.5–9.5 Hz) showed better consolidation of serial-order information. This effect was stronger for more distant channels, highlighting the role of long-range centro-parietal networks underlying the consolidation of serial-order information. The consolidation of probability information was associated with learning-induced changes in delta frequency connectivity (2.5–3 Hz) and seemed to be dependent on more local, short-range connections. Beyond these associations with learning-induced changes, we also found substantial overlap between the baseline and post-learning similarity and their associations with consolidation performance, indicating that stable (trait-like) differences in functional connectivity networks may also be crucial for memory consolidation.

**Significance statement:** We studied memory consolidation in humans by characterizing how similarity in neural oscillatory patterns during learning and rest periods supports consolidation. Previous studies on similarity focused on learning-induced changes (including reactivation) and neglected the stable individual characteristics that are present over resting periods and learning. Moreover, learning-induced changes are predominantly studied invasively in rodents or with neuroimaging or event-related electrophysiology techniques in humans. Here, we introduced a novel approach that enabled us 1) to reveal both learning-induced changes and trait-like individual differences in brain activity and 2) to study learning-induced changes in humans by analyzing continuous EEG. We investigated the consolidation of two types of information and revealed distinct learning-induced changes and trait-like characteristics underlying the different memory processes.

## Introduction

What makes us remember? Successful long-term memory performance depends on consolidation: a process that stabilizes encoded memory representations (McGaugh, 2000). For consolidation, the behavioral, cognitive, and neural states in the first few hours following the learning episode are critical (McGaugh and Izquierdo, 2000). One mechanism that takes place in this crucial time period is the reactivation of the brain activity related to the learning episode (Peigneux et al., 2006; Hermans et al., 2017; Tambini et al., 2017). It has been shown that patterns of brain activity that reappear during the off-line period after learning predict memory consolidation (for a review, see Rasch and Born, 2007). However, previous studies neglected the stable individual characteristics that are present over resting and learning periods and could be relevant for memory consolidation. Here, we aimed to fill this gap by examining how learning-induced changes (including reactivation) and trait-like individual characteristics of neural patterns captured by EEG support memory consolidation.

Post-learning reactivation of memory traces has been widely studied in rodents using invasive neurophysiological techniques (for example, Skaggs and McNaughton, 1996; Nádasdy et al., 1999; Louie and Wilson, 2001). In humans, spontaneous reactivation is mainly studied by non-invasive neuroimaging techniques (PET, fMRI) and it is usually defined by higher similarity of the brain activity patterns during learning and post-learning compared to pre-learning periods (Maquet et al., 2000; Peigneux et al., 2004; Tambini and Davachi, 2019). However, these neuroimaging techniques lack the temporal precision of neuronal activity and are not suitable to capture inter-regional communication through phase synchronization. Examining phase synchronization can shed light on the temporal brain dynamics constituting functional networks underlying consolidation, as oscillatory synchronization integrates anatomically distributed processing and facilitates neuronal communication (Buzsáki and Draguhn, 2004). Furthermore, animal studies have shown that reactivation occurs also in the cerebral cortex, suggesting that it can be studied with scalp EEG (Qin et al., 1997; Ribeiro et al., 2004; Peyrache et al., 2009; Rothschild et al., 2017). Yet, EEG studies of reactivation during post-learning awake rest periods (also termed reinstatement) in humans are still scarce.

Thus, in the current study, we investigated reactivation by comparing functional networks emerging from phase synchronization measured via scalp EEG (Varela et al., 2001) in healthy young adults. We followed a data-driven approach and did not define frequency ranges or brain areas a priori. Rather, we aimed to reveal patterns in both frequency and topography of functional networks that emerge both during learning and post-learning rest. However, similarities in functional networks during learning and subsequent rest do not exclusively emerge as a function of reactivation (or other learning-induced changes), they could also occur due to individual characteristics in brain activity. To disentangle these learning-induced vs. trait-like similarities, we compared the functional connectivity of learning with both pre- and post-learning rest. If the similarity of learning with the pre-learning rest (and its relation to memory consolidation) is comparable to the similarity of learning with the post-learning rest, the similarities likely emerge due to individual characteristics in brain activity. In contrast, higher similarity of learning and post-learning rest compared to pre-learning rest is indicative of learning-induced changes, possibly reactivation.

We used a procedural memory task with an alternating sequence that enabled us to present two types of regularities (serial-order and probability) within the same visual information stream (Nemeth et al., 2013). This design has higher ecological validity, as in real life, we are exposed to different types of information simultaneously. Examining the consolidation of different information simultaneously enabled us to reveal specific similarities in functional networks that uniquely predict the changes in the respective memory performance.

Our work contributes to the understanding of 1) whether (and how) learning-induced changes in EEG connectivity patterns facilitate simultaneous memory consolidation of serial-order vs. probability information, and 2) whether (and to what degree) similarities between learning and post-learning rest and their respective associations with memory consolidation occur due to learning-induced changes (reactivation) or trait-like differences.

## Materials and methods

### Participants

Thirty-four participants (11 males, *M*_age_ = 21.82 ± 2.11) with normal or corrected-to-normal vision were included in the study. Participants were selected from a pool of undergraduate students from Eötvös Loránd University in Budapest. The selection procedure was based on the completion of an online questionnaire assessing mental and physical health status. Respondents reporting no current or prior chronic somatic, psychiatric or neurological disorders or regular consumption of drugs (other than contraceptives) were selected. In addition, individuals reporting occurrences of any kind of extreme life event (e.g., accident) during the last three months that might have had an impact on their mood, affect, and daily rhythms were excluded.

Individuals falling asleep during the post-learning quiet rest (n = 2) were excluded from the analyses. Furthermore, one additional participant was excluded based on extreme behavioral performance. Therefore, the final sample consisted of 31 participants (9 males, *M*_age_ = 21.81 ± 2.10). All participants provided written informed consent before enrollment and received course credits for taking part in the experiment. The study was approved by the research ethics committee of the Eötvös Loránd University, Budapest, Hungary (201410) and was conducted in accordance with the Declaration of Helsinki.

### Task

Behavioral performance was measured by the cued version of the Alternating Serial Reaction Time (ASRT) task (Fig. 1, Nemeth et al., 2013). In this task, a stimulus (picture of a dog’s head or a penguin) appeared in one of four horizontally arranged empty circles on the screen, and participants had to respond by pressing the corresponding button of a response box. Participants were instructed to respond as fast and accurately as they could. The task was presented in blocks with 85 stimuli. A block started with five random stimuli for practice purposes, followed by an 8-element alternating sequence that was repeated ten times. The alternating sequence was composed of fixed sequence (pattern) and random elements (e.g., 2-R-4-R-3-R-1-R, where each number represents one of the four circles on the screen and “R” represents a randomly selected circle out of the four possible ones, Fig. 1B). The response-to-stimulus interval was set to 120 ms. At the end of each block, participants received feedback on their overall accuracy and reaction time and had a 10-20 s rest before starting a new block. As typical in the cued ASRT task, participants were informed about the underlying structure, and their attention was drawn to the alternation of sequence and random elements by different visual cues. In our case, a picture of a dog always corresponded to pattern elements, and a picture of a penguin indicated random elements (Fig. 1A). Participants were instructed to find the hidden pattern defined by the dog in order to improve their performance. Sequence reports (of the pattern elements) were collected at the end of each block after the feedback period. Participants had to type in the regularities they noticed during the task using the same response buttons they used during the ASRT blocks. By the end of the task, all participants reported the correct sequence. On average, participants gained explicit knowledge of the sequence after the 4^th^ block (*M* = 4.10, *SD* = 5.95), and 84% of participants reported the correct sequence during the first 5 blocks.

**Figure 1.**
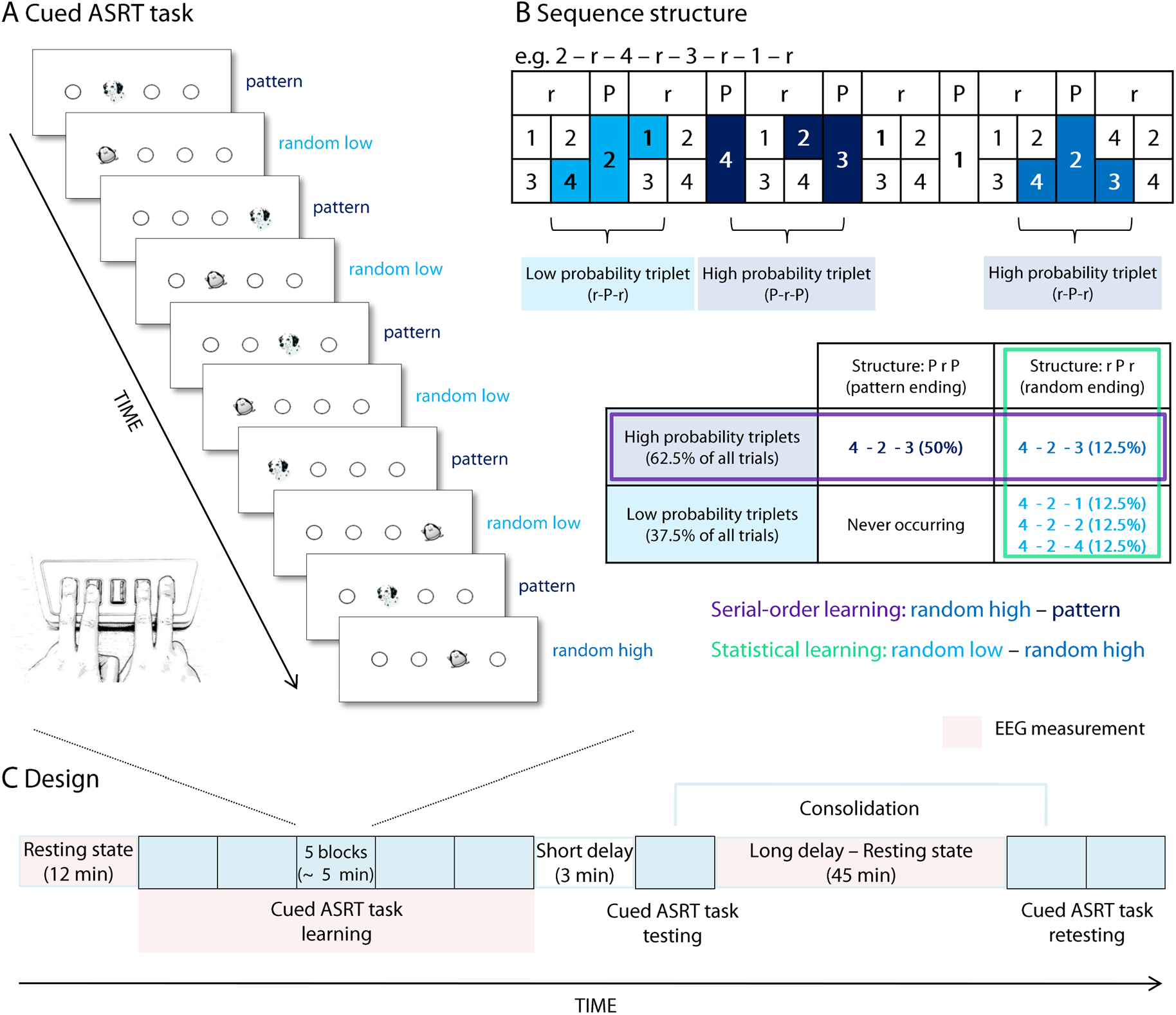
The cued version of the Alternating Serial Reaction Time (ASRT) task. **A)** In this task, pattern and random trials were alternating. Pattern trials were marked with a picture of a dog, whereas random ones with a picture of a penguin. The location of stimuli during the *pattern* trials always followed a predetermined sequence (e.g., 2-4-3-1 in the given example, where numbers correspond to the four locations on the screen), therefore runs of three consecutive elements (triplets) that ended with pattern elements occurred with high probability. Random trials included 1) trials that appeared in an order that was identical to the sequence, therefore they constituted triplets with high probability (*random high* trials) and 2) any other random trials that consequently occurred with low probability (*random low* trials). **B)** We determined for each stimulus whether it was the last element of a high probability triplet (dark blue color) or the last element of a low probability triplet (light blue color) in a sliding window manner. We assessed *serial-order learning* by comparing the responses to pattern trials (which are always high probability triplets) with responses to random high probability trials. *Statistical learning* was quantified as the difference between responses to those random elements that were the last elements of a high probability triplet vs. those that were the last of a low probability triplet. **C)** First, participants were asked to sit quietly with their eyes open and then closed for 6 min each (*pre-learning rest).* Then, the learning period of the ASRT followed that consisted of 25 blocks. To eliminate the fatigue effect and ensure more accurate behavioral consolidation indices, we inserted a short break of 3 min before the testing period of the ASRT that consisted of 5 blocks. After the testing period, there was a 45 min quiet rest period, when participants were sitting with their eyes open and closed in 5 min turns (*post-learning rest).* Finally, there was a retesting period that comprised ten additional ASRT blocks. Behavioral consolidation indices were calculated as the difference in the performance between the beginning of the retesting period and the testing period (5 blocks of each). EEG activity was measured during the pre-learning rest, the learning period of the ASRT and the post-learning rest.

For each participant, one of the six unique permutations of the four possible ASRT sequence stimuli was selected in a pseudo-random manner, so that the six different sequences were used equally often across participants. Note that for the unique permutations, the starting element of a sequence did not matter (e.g., the sequences 2-R-4-R-3-R-1-R and 4-R-3-R-1-R-2-R are identical) because the sequence was continuously presented eight times in a given block.

The task consisted of a total of 40 blocks (Fig. 1C). Participants completed 25 blocks of the ASRT task during the *learning period.* This was followed by a short (3 min) break to minimize the fatigue effect that typically emerges after extended practice (Rickard et al., 2008; Rieth et al., 2010). After the break, participants were tested on the ASRT task for 5 additional blocks that constituted the *testing period.* Subsequently, participants spent an approximately 45 min long off-line period resting quietly. Finally, they completed 10 additional blocks of the ASRT task (*retesting period).* The learning period lasted approximately 30 min, the testing period 5 min, and the retesting period 10 min.

The alternating sequence of the ASRT task formed a sequence structure in which some runs of three consecutive elements (henceforth referred to as triplets) were more probable than others. The more probable triplets were those that formed the sequence, such as 2-X-4, 4-X-3,3-X-1, and 1-X-2 (X indicates any middle element of the triplet) in the above example (2-R-4-R-3-R-1-R). In these triplets, the first and third elements could either be pattern or random stimuli (they constituted 62.5% of all trials). However, the triplets 4-X-1, 4-X-2 or 4-X-4 occurred less likely since the first and third elements could only be random stimuli (each of these triplets occurred in 12.5% of all trials, altogether 37.5%). Figure 1B illustrates this phenomenon with the triplet 4-2-3 occurring with a higher probability than other triplets such as 4-2-1, 4-2-2, and 4-2-4. The former triplet types were termed as high probability triplets, whereas the latter types were termed as low probability triplets (see Fig. 1B and Nemeth et al., 2013). Overall, as a result of the ASRT sequence structure, on the level of individual triplets, the high probability triplets were five times more predictable than the low probability ones.

To calculate learning indices, each stimulus was categorized as either the third element of a high or a low probability triplet. Note that in this way, we determined the probability of each triplet throughout the task in a sliding window manner (i.e., one stimulus was the last element of a triplet, but also the middle and the first element of the consecutive triplets). Moreover, trials were differentiated by the cues (dog and penguin) indicating whether the stimulus belonged to the pattern or random elements. In this way, the task consisted of three trial types: (1) elements that belonged to the predetermined sequence and therefore appeared as the last element of a high probability triplet were called *pattern* trials; (2) random elements that appeared as the last element of a high probability triplet were called *random high* trials; and (3) any other random elements that consequently appeared as the last element of a low probability triplet were *random low* trials (see the example in Fig. 1B).

To disentangle the two key learning processes underlying performance on the cued ASRT task, we differentiated serial-order learning and statistical learning (Fig. 1B). Serial-order learning was measured by the difference in reaction times (RTs) between random high and pattern trials (that is, the average RTs for random high trials minus the average RTs for pattern trials). These trials shared the same statistical properties (both corresponded to the third element of high probability triplets) but had different sequential properties (i.e., pattern vs. random element). Thus, greater serial-order learning was determined as faster responses to pattern vs. random high trials. Note that in previous ASRT studies, serial-order learning was often referred to as sequence learning (Nemeth et al., 2013; Simor et al., 2019). Statistical learning was measured by the difference in RTs between random low and random high trials (that is, the average RTs for random low trials minus the average RTs for random high trials). These trials shared the same sequential properties (both are random) but differed in statistical properties (i.e., they corresponded to the third element of a high or a low probability triplet). Thus, greater statistical learning was determined as faster responses to random high compared to random low trials. In sum, serial-order learning quantified the acquisition of the deterministic rule of the sequential pattern, whereas statistical learning captured purely probability-based learning (Nemeth et al., 2013; Maheu et al., 2020).

### Experimental Design

On the day of the experiment, participants arrived at the laboratory at 10.00 AM. First, an EEG cap with 64 electrodes was fitted by two assistants with impedances set under 10 kΩ. Then, participants were asked to sit quietly with their eyes open and then closed for 6 min each as a baseline EEG recording (*pre-learning rest,* Fig. 1C). Testing with the ASRT started around 11.30 AM and took place in a quiet room equipped with a computer screen, a response box and an EEG recording device. After listening to the instructions, participants had the opportunity to shortly practice the task to get familiar with the stimuli and the response box; however, all stimuli appeared randomly during the practice period. This was followed by the *learning period* of the cued ASRT task that consisted of 25 blocks. To eliminate the effect of fatigue that may have been built up during performing the task for about 25 min (Rickard et al., 2008; Rieth et al., 2010) and therefore ensure more accurate behavioral consolidation indices, we inserted a short break of 3 min during which the fitting of the EEG cap was monitored and impedances were reset under 10 kΩ. After this short delay, the *testing period* of the ASRT followed that consisted of 5 blocks. After the testing period, we introduced a long quiet rest period, during which participants were sitting in a dim lit room, facing towards an empty wall for 45 min (*post-learning rest*). During this resting state period, to prevent participants from falling asleep, we instructed them to open and close their eyes periodically (approximately every 5 min). Furthermore, the EEG activity was monitored online, and if signs of sleepiness (alpha disappearance and theta increase) occurred, participants were instructed to open their eyes and/or make movements. After the resting period, we set the impedances again under 10 kΩ. Finally, there was a *retesting period* that comprised 10 ASRT blocks. Behavioral consolidation indices were calculated as the difference in performance between the beginning of the retesting period and the testing period (5 blocks each). EEG activity was measured during the pre-learning rest, the learning period of the ASRT and the post-learning rest.

### EEG recording

EEG activity was measured using a 64-channel recording system (BrainAmp amplifier and BrainVision Recorder software, BrainProducts GmbH, Gilching, Germany). The Ag/AgCl sintered ring electrodes were mounted in an electrode cap (EasyCap GmbH, Herrsching, Germany) on the scalp according to the 10% equidistant system. During acquisition, electrodes were referenced to a scalp electrode placed between the Fz and Cz electrodes. Horizontal and vertical eye movements were monitored by EOG channels. Three EMG electrodes to record muscle activity, and one ECG electrode to record cardiac activity were placed on the chin and chest, respectively. All electrode contact impedances were kept below 10 kΩ. EEG data were recorded at a sampling rate of 500 Hz, band-pass filtered between 0.3 and 70 Hz.

### Data analysis

#### Behavioral data

For the analysis of the cued ASRT, we followed procedures outlined in previous studies (Howard and Howard, 1997; Song et al., 2007; Nemeth et al., 2013; Simor et al., 2019). First, the blocks of the task were collapsed into epochs of five blocks to facilitate data processing and to increase statistical power. The first epoch contained blocks 1–5, the second epoch contained blocks 6–10, etc. We calculated median reaction times (RTs) for all correct responses, separately for pattern, random high and random low trials, and for each epoch. Two kinds of low probability triplets were eliminated: repetitions of a single element (e.g., 2-2-2, 3-3-3) and triplets beginning and ending with the same element (e.g., 2-1-2, 3-4-3) as individuals often show pre-existing response tendencies to such triplets (Howard et al., 2004). Thus, we eliminated these triplets to ensure that differences between high vs. low probability triplets emerged due to learning and not to pre-existing response tendencies.

### Overall performance trajectory

To evaluate performance changes in RTs due to learning throughout the entire task, we conducted a repeated-measures analysis of variance (ANOVA) on RTs with EPOCH (1–8) and TRIAL TYPE (pattern, random high, random low) as within-subject factors. Post-hoc comparisons were performed using Fisher’s LSD test. Then, we computed a serial-order and a statistical learning score for each epoch. The serial-order learning score was calculated as the difference between the RTs for random high trials minus the RTs for pattern trials. The statistical learning score was calculated as the difference between the RTs for random low trials minus the RTs for random high trials. In both cases, higher learning scores indicated better learning.

### Consolidation

To examine off-line changes occurring between testing and retesting periods, we used a repeated-measures ANOVA on the learning scores, with EPOCH (6–7, the only epoch of the testing period and the first epoch of the retesting period) and LEARNING TYPE (serial-order learning, statistical learning) as within-subject factors. For the ANOVAs, Greenhouse-Geisser epsilon (e) correction (Greenhouse and Geisser, 1959) was used to adjust for a lack of sphericity when necessary. Original df values and corrected p values (if applicable) are reported together with partial eta-squared (η_p_^2^) as the measure of effect size. To be able to identify correlations between brain activity patterns and memory performance, we computed consolidation indices: off-line changes in serial-order and statistical learning were defined as the difference between the learning scores at the beginning of the retesting period (epoch 7) minus the learning scores in the testing period (epoch 6; Fig. 1C). A positive value indicated an improvement in learning performance during the off-line period. For all subsequent analyses between brain activity and memory performance, we used these consolidation indices.

#### EEG data

Preprocessing and further analysis (Fig. 2A) of the EEG data were performed in MATLAB (version R2017b, The Mathworks, Inc, Natick, MA) using the EEGLAB (14_0_0b version, Delorme, Sejnowski, & Makeig, 2004) and the Fieldtrip (Oostenveld et al., 2011) toolboxes. The following steps were executed for EEG data recorded during the pre-learning rest, the learning period of the ASRT and the post-learning rest.

**Figure 2.**
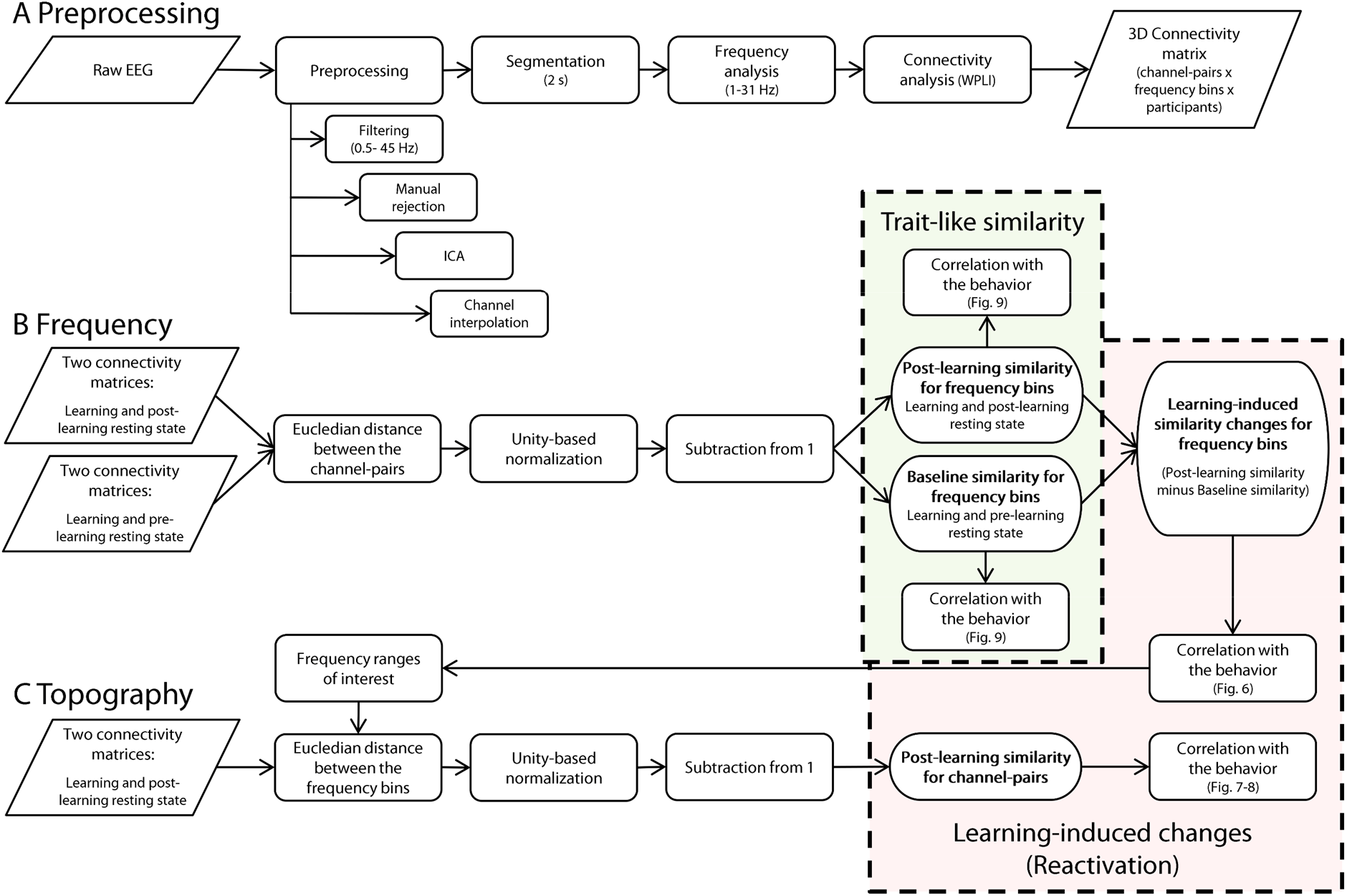
Steps of the analysis of EEG data. **A)** Connectivity matrices were calculated from the raw EEG data for the pre-learning rest, the learning period of the ASRT and the post-learning rest, resulting in 3D connectivity matrices for these three periods with channel-pairs, frequency bins and participants as dimensions. **B)** Comparing two connectivity matrices at a time (learning and post-learning rest or learning and pre-learning rest), we computed the similarity of functional connectivity during these different periods (henceforth referred to as Post-learning and Baseline similarity, respectively) for frequency bins between 1 and 31 Hz. We calculated the Euclidean distance between all channel-pairs for each frequency bin. Then, we normalized these values to shift the scale to the range of 0 to 1 and subtracted them from 1 to indicate similarity rather than distance, with higher values indicating greater similarity. Then, we performed correlational analyses between the consolidation indices and these Post-learning and Baseline similarity values for frequency bins, which could reveal trait-like associations between behavior and similarity of neural patterns. To reveal learning-induced changes (possibly reactivation) in similarity, we subtracted the Baseline similarity values from the Post-learning similarity values, with larger values indicating greater similarity between the learning period and post-learning rest than between the learning period and pre-learning rest. We identified frequency ranges of interest for further analysis based on the correlations that these similarity difference values showed with the consolidation indices for consecutive frequency bins. **C)** As a second step, we computed Euclidean distance between the identified frequency bins of the learning and post-learning rest connectivity matrices, which we transformed again to similarity values by normalization and subtraction from 1. This resulted in Post-learning similarity values for each channel-pair within the frequency ranges of interest. Finally, we contrasted these Post-learning similarity values for channel-pairs with behavior to investigate the topographical patterns of the associations revealed in the frequency domain. With the steps outlined in B) and C), we aimed to differentiate trait-like (light green box) and learning-induced (light orange box) associations between memory consolidation and similarity of functional connectivity across different periods (see main text for details).

For preprocessing, data were off-line band-pass filtered (0.5–45 Hz) using Hamming windowed finite-impulse-response filter. Then, task-free periods of brain activity during the learning period (feedback and sequence typing periods of the task and the resting periods between blocks) and high amplitude artifacts (body movements, sweating, and temporary electrode malfunction) were visually rejected. Subsequently, to identify (horizontal and vertical) eye-movement artifacts, we ran the extended infomax algorithm of independent component analysis (Lee et al., 1999) via EEGLAB (option: runica, ‘extended’). ICA components constituting eye-movement artifacts were removed via visual inspection of their time-series, topographical distribution and frequency contents. Maximum six independent components (out of 56 components produced by the algorithm) per participant were removed. Additionally, if needed, bad channels were interpolated for a maximum of four bad channels per participant.

After artifact rejection, non-overlapping epochs of 2000 ms duration were extracted from the continuous EEG recording. This data segmentation yielded *M* = 319.77, *SD* = 33.30, *M* = 707.94, *SD* = 67.43, *M* = 988.74, *SD* = 269.92 epochs for the pre-learning rest, learning, and post-learning rest, respectively. Due to more body movement artifacts during the post-learning rest, proportionally more data from this period had to be removed. Note that the EEG during the learning period was segmented regardless of the onsets of stimuli (i.e., epochs had random timing relative to trials). To compute power spectral density, Hann taper was first applied to the two-second long, artifact-free EEG segments. Then, for each participant and channel, power spectral density was computed for each 0.5 Hz frequency bin between 1 and 31 Hz with the Fast Fourier Transform (FFT) algorithm as implemented in Fieldtrip.

Functional connectivity was calculated between all channel pairs by measuring phase synchronization using the Weighted Phase Lag Index (WPLI) as implemented in Fieldtrip (ft_connectivityanalysis function, ‘wpli’ option). Phase Lag Index (PLI) was introduced by (Stam et al., 2007), and it reflects the consistency by which one signal is phase leading or phase lagging with respect to another signal. PLI has been shown to be sensitive in detecting dynamic changes of phase relationships between different brain areas, and to be insensitive to the effect of volume conduction (effect of common sources of the EEG signal), as well as to be (largely) independent of the reference electrode. PLI attenuates the volume conduction effect by disregarding phase lags of zero or π (as these phase differences suggest two electrodes are picking up signal from the same source). The WPLI is a modified version of PLI that weights phase angle differences around 0.5 and 1.5 π more than those around zero and π, which makes the method more robust against volume conduction (Vinck et al., 2011). Note that the Fieldtrip function results in WPLI values between −1 and 1, with both −1 and 1 indicating higher synchronization. In our analyses, we used the absolute WPLI values as the measure of the strength of the interaction. Thus, WPLI values in our analyses fall between 0 and 1, with higher values indicating more consistent phase differences and higher synchronization.

The mentioned steps resulted in three 3D connectivity matrices, separately for the pre-learning rest, the learning period of the ASRT and the post-learning rest (Fig. 2A). The dimensions in these matrices were channel-pairs, frequency bins and participants. To measure similarity between the brain activity recorded at different periods, we calculated Euclidean distance between these connectivity matrices (Fig. 2BC).

#### Associations between behavior and similarity in EEG functional connectivity across different periods

To reveal associations between behavioral indices of consolidation and brain activity, we computed similarity between functional connectivity measured during the learning period and the pre-learning rest, and between the learning period and the post-learning rest separately (henceforth referred to as *Baseline* and *Post-learning similarity,* respectively). We computed two types of similarity: 1) for frequency (Fig. 2B) and 2) for channel-pairs (Fig. 2C). As a data-driven approach, we identified frequencies associated with the behavioral indices of consolidation first. Then, we explored relevant topographical patterns in the identified frequency ranges based on the channel-pair similarity.

### Frequency of reactivation

To reveal possible reactivation (reinstatement) of neural patterns after learning, we aimed to find frequencies 1) where the Post-learning similarity is different from the Baseline similarity, presumably reflecting learning-induced changes, and 2) where this similarity difference is linked to memory performance. We calculated the Baseline and Post-learning similarity values for frequency bins from 1 to 31 Hz as follows: We computed Euclidean distance between all channel-pair connectivity values of the learning period and the pre-and post-learning rest respectively, separately for each subject and frequency bin. The formula of the Euclidean distance for frequency bins is:

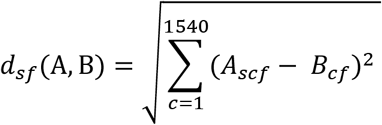

where A and B denote the studied two connectivity matrices and the index variables *s*, *c* and *f* indicate subject, channel-pair and frequency bin, respectively.

Then, we transformed these distance values into similarity values by unity-based normalization (linear transformation to shift the scale to the range of 0 to 1) and subtracted these values from 1. In this way, we obtained a similarity value for each subject and each frequency bin, separately for Baseline and Post-learning similarity (Fig. 2B). The similarity values fell between 0 and 1, where higher values indicated greater similarity. We then calculated the *Learning-induced similarity change* values by subtracting the Baseline similarity values from the Post-learning similarity values, where higher values indicated greater similarity between learning and post-learning rest compared to learning and pre-learning rest. These Learning-induced similarity change values reflected the changes in the resting state functional connectivity values that appear after the learning episode compared to the pre-learning resting state. Finally, to detect associations with behavioral measures, we computed Spearman correlations between the Learning-induced similarity change values and the two consolidation indices (consolidation of serial-order and statistical knowledge) for each frequency bin (Fig. 2B). We identified candidate oscillatory frequencies based on clusters of neighboring frequency bins showing similar, significant associations with the same consolidation index.

### Controlling for multiple comparisons

To account for the problem of multiple comparisons, we ran a nonparametric cluster-based permutation correction for the correlations between the consolidation indices and the Learning-induced similarity change values (Cohen, 2014). Our aim was to find neighboring frequency bins (frequency ranges) that showed similar associations between behavior and brain activity, indicating plausible oscillatory activity relevant for consolidation. The emergence of clusters of frequency bins that display similar associations can indicate true effects. Thus, we accounted for the multiple comparison problem on the cluster level. We created test statistics under the null hypothesis of no association by randomly shuffling the behavioral index for 1000 iterations and calculating correlations between these shuffled behavioral and the original Learning-induced similarity change values. We then created clusters for the cluster-based correction by thresholding these iteration maps by a p-value based on the standardized Z-values of null-hypothesis test statistics (pre-cluster threshold). Then we used the Matlab function ‘bwconncomp’ to identify clusters in each thresholded iteration map. This yielded a distribution of the largest pre-cluster suprathreshold clusters that can be expected under the null hypothesis. As a final step, we compared the suprathreshold clusters in the original statistical values and removed any clusters that were less than the 95 percentile of the distribution of the largest clusters under the null hypothesis. With this method, we identified frequency ranges in which greater Learning-induced similarity change values (that is, greater Post-learning similarity compared to Baseline similarity) showed robust associations with behavior.

### Topography of reactivation

To reveal topographical information of the functional connections in the identified frequency ranges of interest, we computed similarity values for channel-pairs (Fig. 2C). The frequency ranges of interest were identified based on the correlations between the behavioral consolidation indices and the Learning-induced similarity change values (see above). For these frequency ranges of interest, we calculated Euclidean distance between all relevant frequency bin connectivity values in the learning period and the post-learning rest, for each subject and channel-pair separately. The formula of the Euclidean distance for channel-pairs is:

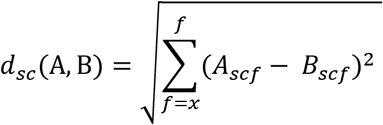

where A and B denote the studied two connectivity matrices and the index variables *s, c* and *f* indicate subject, channel-pair and frequency bin, respectively. As there are multiple frequency ranges where we compute the distance for the channel-pairs, x denotes the lowest frequency in each of these frequency ranges.

Again, to yield similarity values, we used unity-based normalization and subtracted the Euclidean distance values from 1. To obtain the topography of functional connections relevant for memory consolidation within the frequency ranges of interest, we computed Spearman correlations separately for each channel-pair similarity value and the two consolidation indices. Note that for these channel-pair correlations, we used the Post-learning similarity rather than the Learning-induced similarity change values because we computed the channel-pair similarity only in the frequency ranges where this change has been accounted for. Also, as our aim was to identify topographical patterns in functional connections (rather than identifying specific connections more reliably), we did not correct for multiple comparisons in these correlation analyses. For the same purpose, we differentiated positive and negative correlations in all analyses that targeted the exploration of topographical patterns in relevant functional connections.

### Patterns in topography

To explore patterns in the topography of functional connections in the relevant frequency ranges, we first aimed to investigate how the range of the connections (short- or long-range) is associated with consolidation performance. We analyzed the associations between the distances of the electrodes in the channel-pairs and the correlation coefficients that were obtained between the Post-learning similarity values of those respective channel-pairs and the consolidation indices. Importantly, we investigated these associations separately for the consolidation of serial-order vs. statistical knowledge, for those respective frequency ranges where they showed significant associations with the EEG similarity values. To evaluate whether the distance between the electrodes in a channel-pair influenced the associations, we conducted one-way ANOVAs on the relevant Spearman correlation coefficients with topographical DISTANCE of channel-pairs (short: first 25% of distances (0-25%), medium: middle 25% of distances (37.5-62.5%), long: last 25% of distances (75-100%)). To include all (positive and negative) associations in this analysis, we used the absolute values of the correlation coefficients. To evaluate the effect of distance in more detail, we performed pairwise comparisons using Fisher’s LSD test. Welch’s ANOVA and Games-Howell post-hoc comparisons were used if homogeneity of variances was violated. Next, we aimed to investigate whether differences in associations due to distance are present both for positive and negative correlations. Therefore, we differentiated between positive and negative correlations and fitted a linear trendline for these associations, separately for serial-order and statistical consolidation in the relevant frequency ranges identified in the previous analyses.

### Trait-like similarity

Finally, to contrast trait-like and learning-induced similarities, we compared the Baseline and the Post-learning similarity values for frequency bins and their associations with behavioral indices of consolidation. We computed Spearman correlations (similarly to the correlations described in the ‘Frequency of reactivation’ section) between the consolidation indices and the ‘raw’ Baseline and Post-learning similarity values for the frequency bins between 1 and 31 Hz. For these analyses, we again ran a nonparametric cluster-based permutation correction (described in the ‘Controlling for multiple comparisons’ section).

### Controlling for the influence of the raw functional connectivity

It is possible that the correlations emerging between the similarity/learning-induced similarity change values for frequency bins and the consolidation indices were dominated by the functional connectivity of one of the periods of learning or pre-/post-learning rest. To exclude this possibility, we again computed Spearman correlations (similarly to the correlations described in the ‘Frequency of reactivation’ section) between the consolidation indices and the ‘raw’ functional connectivity measures averaged for the frequency bins between 1 and 31 Hz separately for the pre-learning rest, learning and post-learning rest. For these analyses, we again ran a nonparametric cluster-based permutation correction (described in the ‘Controlling for multiple comparisons’ section).

## Results

### Behavioral data

#### Overall performance trajectory

The ANOVA on the overall performance trajectory in the ASRT task showed that irrespective of trial type, RTs significantly decreased across epochs (main effect of EPOCH: *F*_7,210_ = 80.748, *p* < .0001, η_p_^2^ = .73), indicating increasingly faster RTs due to practice (Fig. 3A). Furthermore, participants showed significant serial-order and statistical learning (main effect of TRIAL TYPE: *F*_2,60_ = 25.065, *p* < .0001, η_p_^2^ = .45): they responded faster to pattern compared to random high trials (*p* < .0001), and faster to random high compared to random low trials (*p* < .0001). The EPOCH x TRIAL TYPE interaction was also significant (*F*_14,420_ = 4.689, *p* = .003, η_p_^2^ = .14), indicating different learning trajectories for the three trial types (see Fig. 3A). Although participants became faster for all trial types during the course of the task, responses to pattern trials showed greater gains in comparison to both random high and random low trials: Average RTs of pattern trials decreased from 355.65 to 253.98 ms (*p* < .0001), of random high trials from 376.95 to 320.52 ms (*p* < .0001), and of random low trials from 392.07 to 349.87 ms (*p* < .0001). The speed-up in responses to pattern trials was significantly larger than that to random high trials (*t*_30_ = 2.72, *p* = .011), indicating increasing serial-order learning as the task progressed. The speed-up in responses to random high compared to random low trials was also significantly larger (*t*_30_ = 2.07, *p* = .047), indicating increasing statistical learning as the task progressed.

**Figure 3.**
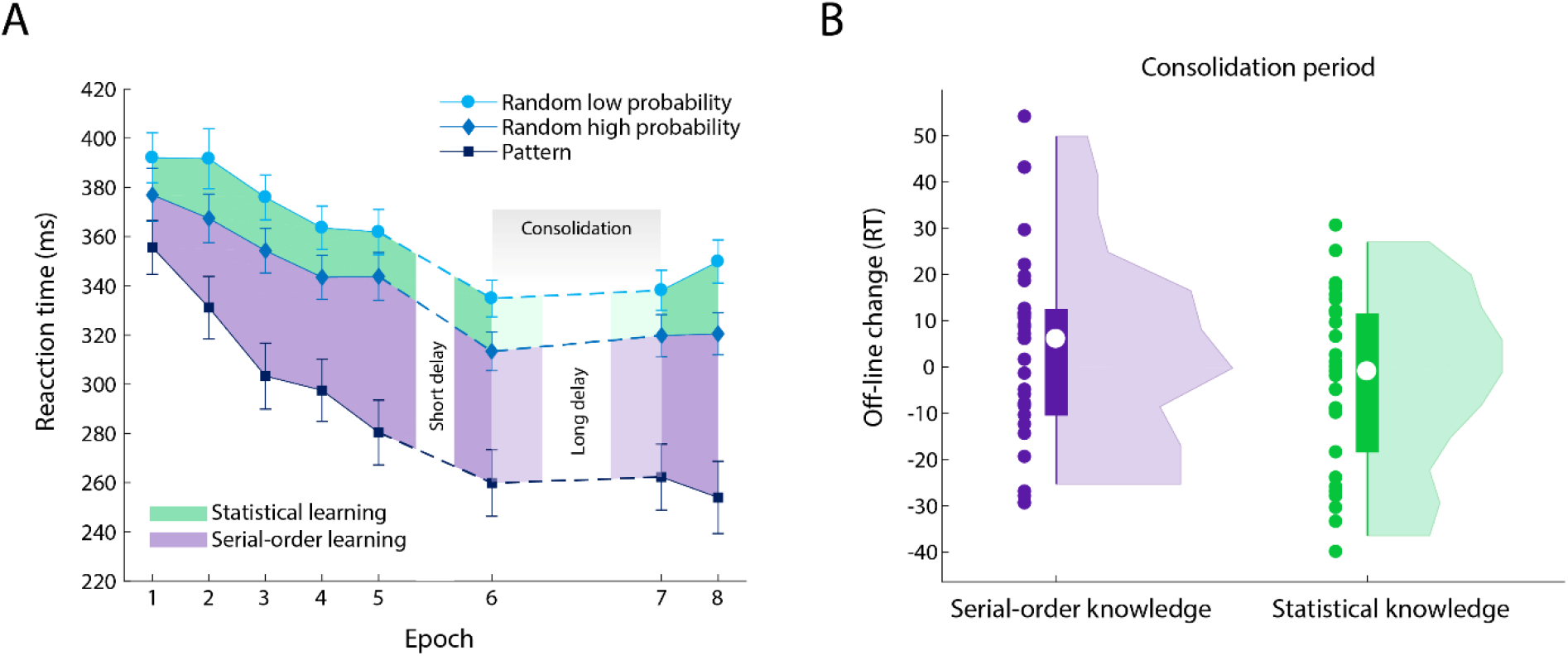
Performance in the ASRT task. **A)** Performance during the learning (epochs 1–5), testing (epoch 6) and retesting (epochs 7–8) periods. Mean reaction times (RTs) across participants are shown for pattern (dark blue line with square symbols), random high (medium blue line with rhombus symbols), and random low (light blue line with circle symbols) trials in each epoch. Error bars denote standard error of the mean (SEM). The difference of RTs for pattern and random high trials indicates serial-order learning (purple shading). The difference in RTs for random high and random low trials indicates statistical learning (green shading). **B)** Off-line changes in serial-order knowledge (purple) and statistical knowledge (green). Off-line changes were calculated as the learning scores of epoch 7 minus the respective learning scores of epoch 6. The plot shows rotated probability density of these indices, the dots indicate individual data points, the bars represent data between the first and third quartiles, and the white circles within the bars show the medians.

#### Consolidation of serial-order and statistical knowledge

To evaluate the changes in the memory performance during the off-line period that followed the learning, we computed learning scores for serial-order and statistical knowledge separately (for details see ‘Behavioral data’ section in Data analysis). The ANOVA to evaluate the off-line changes in serial-order and statistical knowledge showed that participants exhibited overall higher scores in serial-order than in statistical knowledge (main effect of LEARNING TYPE: *F*_1,30_ = 6.284, *p* = .018, η_p_^2^ = .17). However, overall learning scores (regardless of learning type) were stable from the testing period to the first retesting epoch (main effect of EPOCH: *F*_1,30_ = 0.064, *p* = .802, η_p_^2^ = .002). Furthermore, the off-line changes (consolidation) of serial-order and statistical knowledge did not appear to be different (EPOCH x LEARNING TYPE interaction: *F*_1,30_ = 1.356, *p* = .253, η_p_^2^ = .043), suggesting retention of both types of knowledge in the offline period. For further analyses, we computed consolidation indices separately for serial-order and statistical knowledge (see ‘Behavioral data’ section in Data analysis). Importantly, the variance of the consolidation indices appeared to be sufficient for subsequent analyses investigating associations between brain activity and memory consolidation performance (Fig. 3B).

### EEG data

Both spectral power and functional connectivity analyses showed higher synchrony in the alpha frequency in the pre- and post-learning rest periods compared to the learning period (Fig. 4).

**Figure 4.**
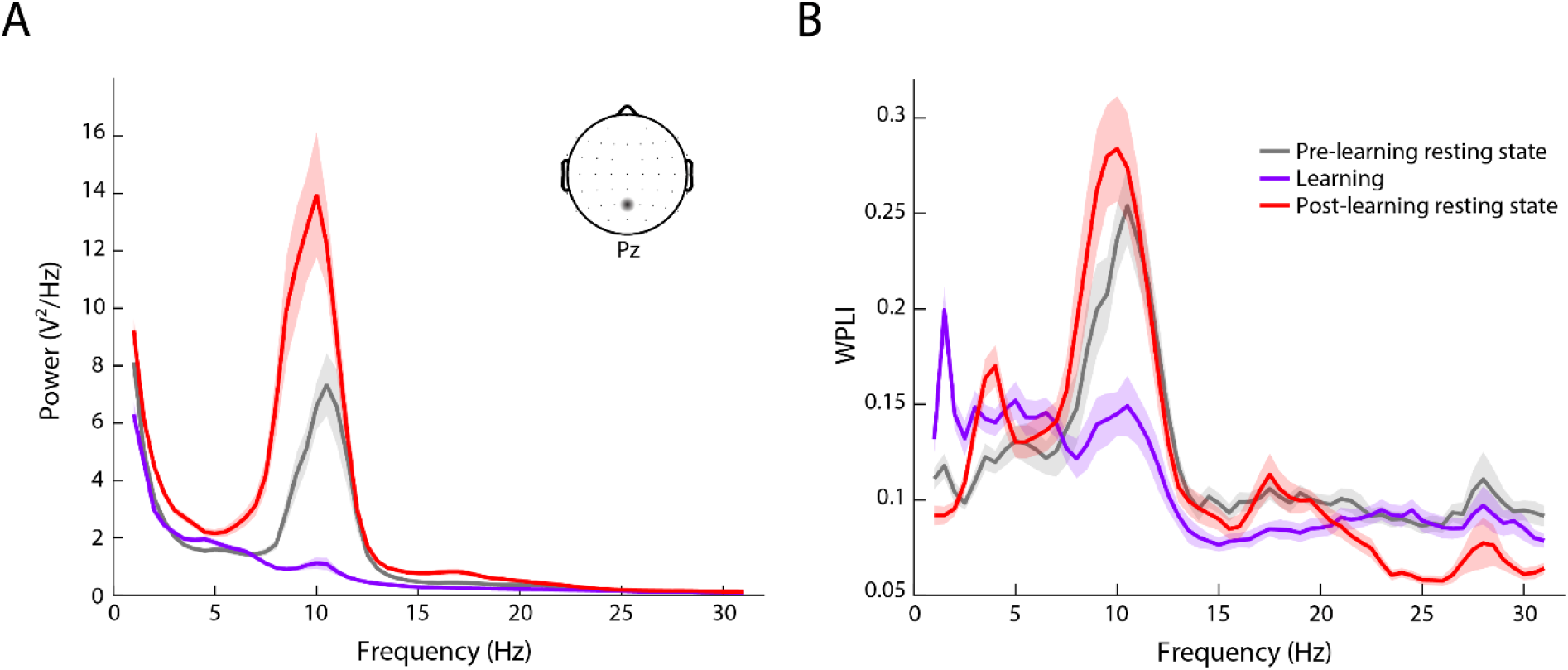
**A)** Example of spectral power in one channel (Pz) and **B)** average connectivity (across all channels) measured by the Weighted Phase Lag Index for frequency bins between 1 and 31 Hz for the pre-learning rest (gray), ASRT learning (purple), and post-learning rest (red) periods. The solid lines indicate the mean and the shaded error bars indicate SEM of values averaged across all participants.

### Associations between behavior and similarity in EEG functional connectivity across different periods

#### Frequency-wise similarity analysis

We computed similarity between the functional connectivity of the learning period and pre-learning rest (Baseline similarity) and the learning period and post-learning rest (Post-learning similarity) for frequencies 1 to 31 Hz (see Fig. 2 and related text in Data analysis). These Baseline and Post-learning similarity values computed for frequency bins exhibited similar patterns (Fig. 5A): They both showed on average high similarity (closer to the maximum value of 1) and a steep fall in similarity in the alpha frequency range. This drop in the similarities in the alpha frequency is expected, given the marked differences between alpha synchronization (both in spectral power and phase-synchronization, Fig. 4) during learning and the pre-/post-learning resting states. To reveal changes in the resting state functional connectivity values that appear after the learning episode, we calculated Learning-induced similarity change values (the difference between the Baseline and Post-learning similarities, see also Fig. 2B and related text). While the averages of the Learning-induced similarity change values across participants were around 0, the individual differences (Fig. 5B) enabled us to compute correlations with the behavioral indices of memory consolidation. As we aimed to reveal associations between memory consolidation and brain activity, we focused on similarities that correlated with behavior, and not on the emergence of similarities per se.

**Figure 5.**
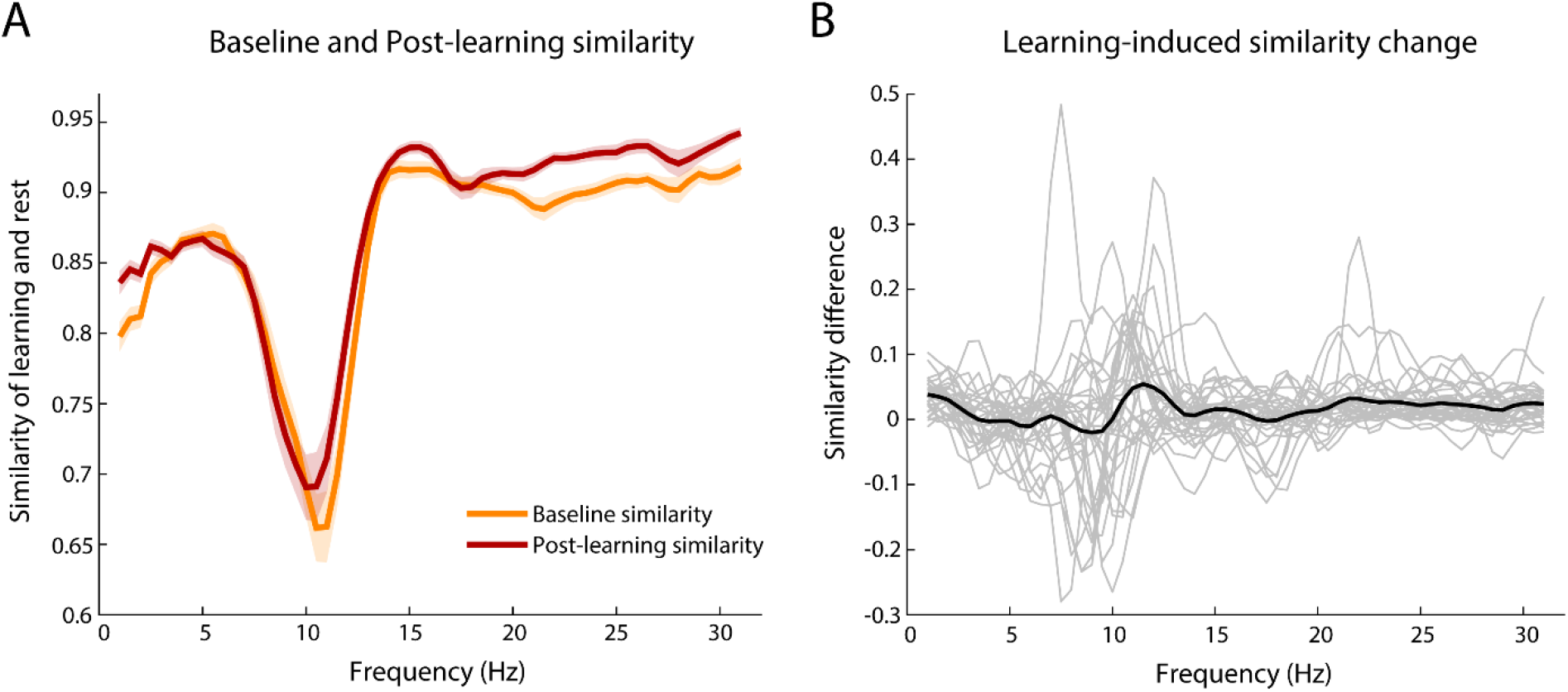
**A)** Similarity between connectivity matrices of the learning period and the pre-learning rest (Baseline similarity), and the learning period and the post-learning rest (Post-learning similarity) for frequency bins. Baseline and Post-learning similarity exhibited similar patterns. The solid lines indicate the mean and the shaded error bars indicate the SEM for Baseline (orange) and Post-learning (red) similarity. **B)** Learning-induced similarity change was calculated as the Post-learning similarity values minus Baseline similarity values, with larger values indicating greater similarity between the learning period and post-learning rest than between the learning period and pre-learning rest. Thinner gray lines represent each participant, while the thicker black line indicates the mean averaged across all participants.

#### Frequency of reactivation

To identify the oscillatory frequencies where the Learning-induced similarity change values were associated with the behavioral indices of consolidation, we computed correlation coefficients (Fig. 6A). Consolidation of serial-order knowledge showed a significant positive correlation with the Learning-induced similarity change in the alpha frequency range (for the 8.5-9.5 Hz bins, Fig. 6B). In contrast, consolidation of statistical knowledge was positively associated with the Learning-induced similarity change in the delta frequency (for the 2.5 and 3 Hz bins, Fig. 6C). This indicates that the more similar the functional connections were between the learning period and post-learning rest compared to the learning period and the pre-learning rest in the alpha and delta frequencies, the better the consolidation was for serial-order and statistical knowledge, respectively.

**Figure 6.**
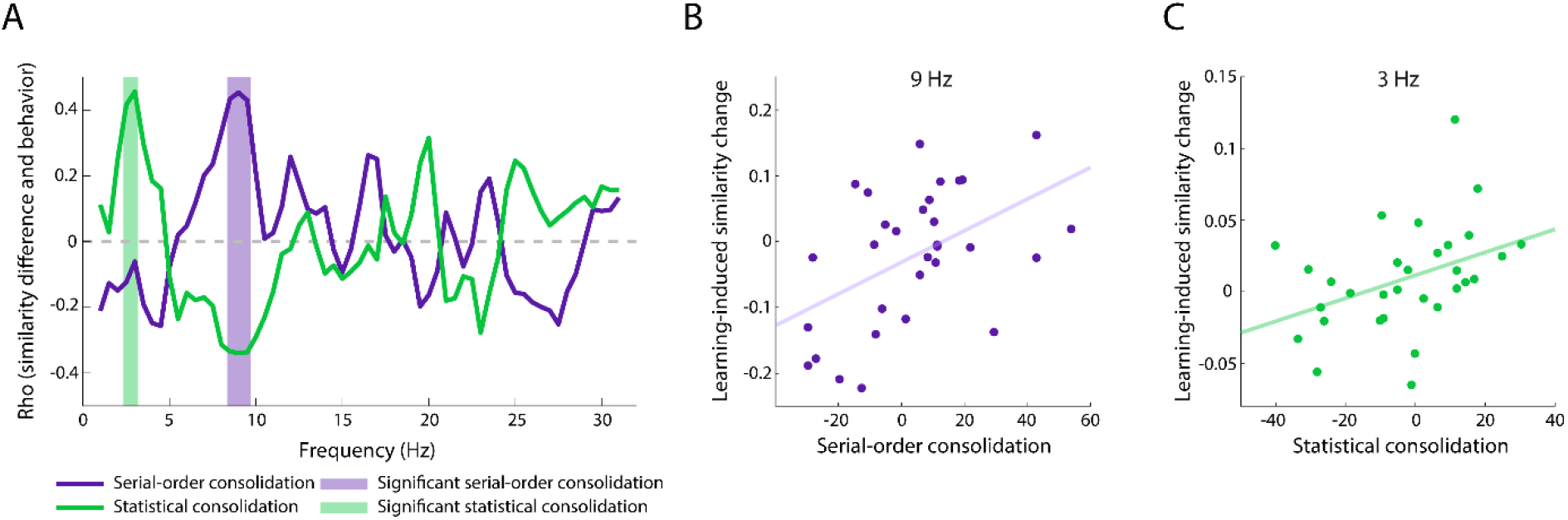
**A)** Correlation coefficients (Spearman Rho’s) between the behavioral indices of consolidation and the Learning-induced similarity change values (difference between Baseline and Post-learning similarity). Purple and green lines indicate associations with serial-order and statistical consolidation, respectively. The shading denotes the significant (*p* < .05) correlations after cluster-based correction for multiple comparisons. **B)** Positive **c**orrelation between the Learning-induced similarity change value at 9 Hz and the consolidation of serial-order knowledge. **C)** Positive correlation between the Learning-induced similarity change value at 3 Hz and the consolidation of statistical knowledge. On B) and C), the dots represent individual data points for each participant and the lines are linear trendlines.

#### Topography of reactivation

Next, we aimed to reveal patterns in the topography of functional connections that were present both during learning and post-learning rest, and could indicate reactivation (see Fig. 2C and related text). We computed channel-pair similarities within the frequency ranges (alpha and delta) where the Learning-induced similarity change values showed significant associations with the consolidation indices. As we showed that higher Post-learning similarity compared to Baseline similarity in these frequency ranges is beneficial for memory consolidation, we can assume that Post-learning channel-pair similarities in these frequency ranges are indicative of reactivation or other learning-induced changes. To find relevant channel-pairs where the similarity is associated with the consolidation indices, we computed correlations between the Post-learning channel-pair similarities within the identified frequency ranges and the respective consolidation index: channel-pair similarity in the significant alpha frequency range (8.5-9.5 Hz) with the consolidation of serial-order knowledge, and channel-pair similarity in the significant delta frequency range (2.5-3 Hz) with the consolidation of statistical knowledge. Consolidation of serial-order knowledge showed significant positive correlations with channel-pair similarity in the alpha frequency over the whole brain, particularly at centro-parietal channel pairs (Fig. 7A). In contrast, for the consolidation of statistical knowledge, mainly significant negative correlations emerged with channel-pair similarity in the delta frequency over the whole brain, particularly over posterior and lateral sites (Fig. 7B).

**Figure 7.**
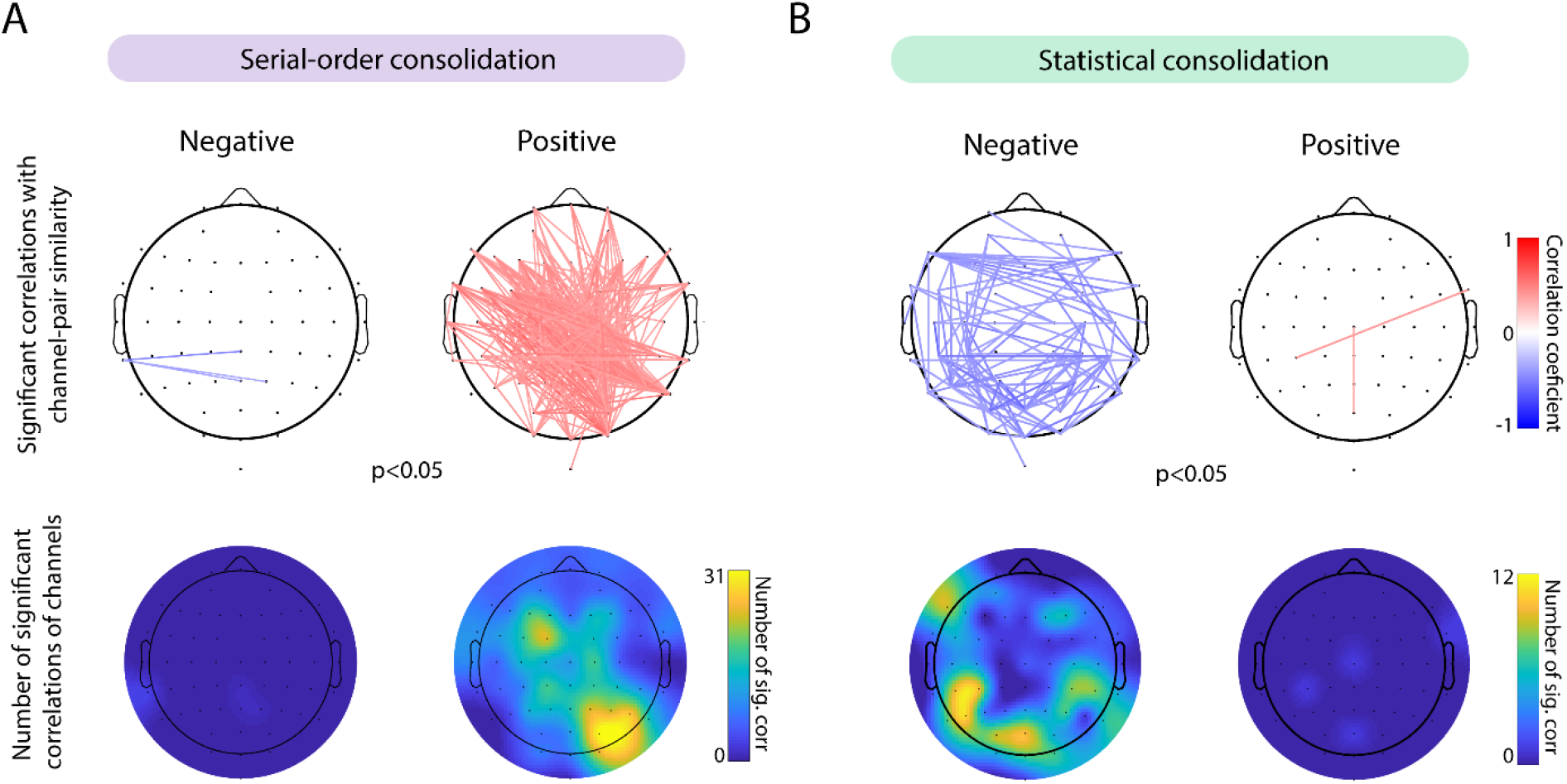
**A)** Topography of significant correlations between the consolidation of serial-order knowledge and Post-learning similarity for channel-pairs in the (8.5-9.5 Hz) alpha frequency range. **B)** Topography of significant correlations between the consolidation of statistical knowledge and Post-learning similarity for channel-pairs in the (2.5-3 Hz) delta frequency range. In the upper panel, significant (uncorrected) positive and negative correlations are shown. In the lower panel, for each channel, the number of significant correlations in which the given channel appears is shown separately for positive and negative associations.

#### Patterns in topography

Next, we aimed to explore patterns in the correlation of all channel-pair similarities with memory consolidation. More specifically, we investigated whether the strength of the associations between the channel-pair similarities and the consolidation indices vary as a function of the distance (short-, medium-vs. long-range) of functional connections. Importantly, we investigated these associations separately for the consolidation of serial-order and statistical knowledge, for the respective frequency ranges where they showed significant associations with the similarity values. Note that we included all correlation coefficients in subsequent analyses, not only the significant ones depicted in Figure 7.

The ANOVA on the correlation coefficients between the consolidation index of serial-order knowledge and the channel-pair similarity values in the alpha frequency range as a function of the distance between the electrodes of the channel-pairs was significant (F_2,1149_ = 28.00, *p* < .001, η_p_^2^ = .046): absolute values of correlation coefficients were different as a function of distance (Fig. 8A, left). Post-hoc tests revealed that channel-pairs with medium or long distances showed significantly stronger correlations with serial-order consolidation than channel-pairs with short distances (*p*s < .001), while correlation strengths did not differ between medium and long distances (*p* = .14). When considering the direction of correlations (positive or negative correlation coefficients), we found that the trend visible in the ANOVA, namely that the consolidation of serial-order knowledge showed stronger associations with longer connections was led by the positive correlation coefficients (Fig. 8A, right).

**Figure 8.**
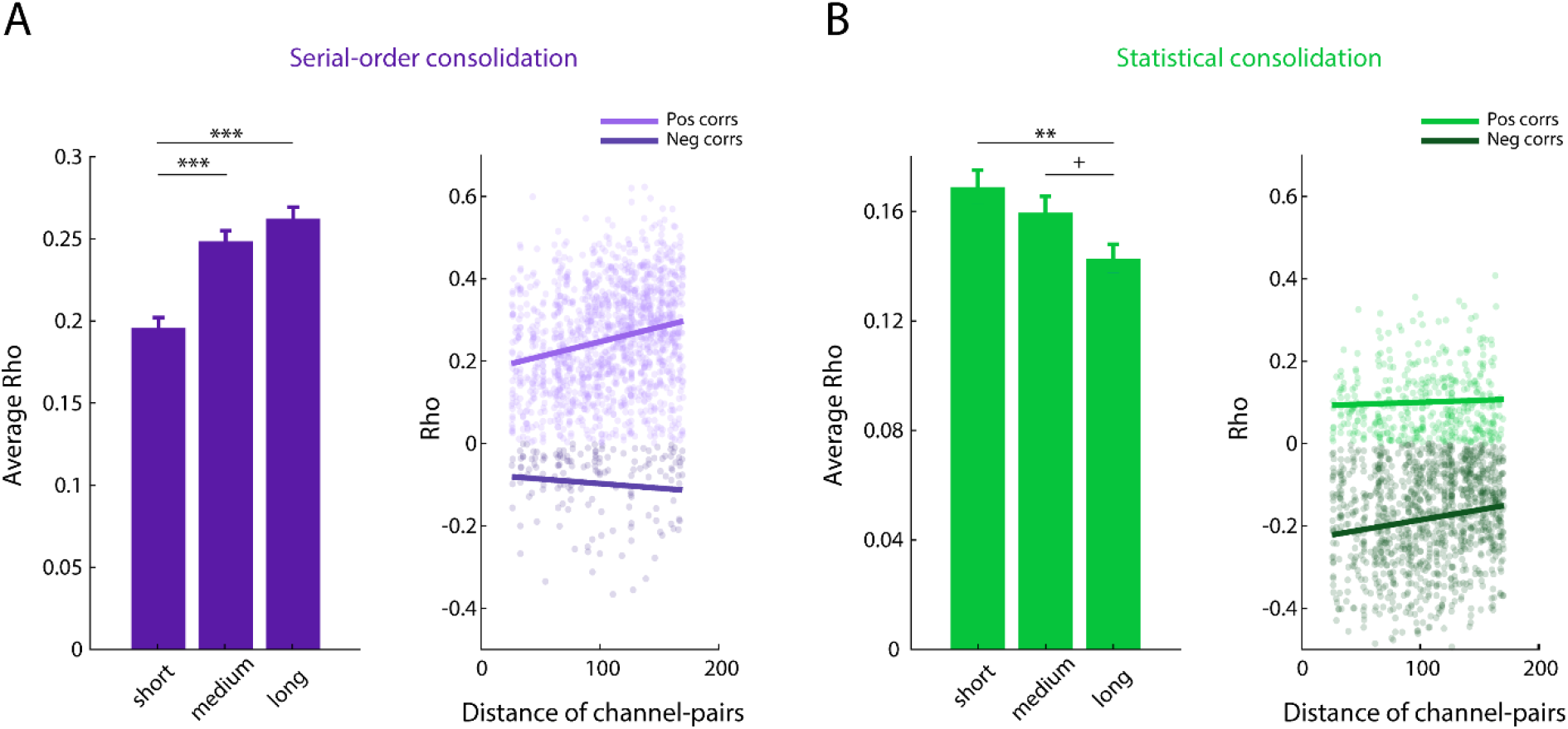
Associations between the Post-learning channel-pair similarity values and the behavioral indices of consolidation as a function of the distance between the electrodes of the channel-pairs. **A)** For the consolidation of serial-order knowledge (purple), associations in the alpha frequency range are shown. These associations between Post-learning channel-pair similarity in the alpha frequency and serial-order consolidation were more pronounced in more distant channel-pairs. **B)** For the consolidation of statistical knowledge (green), associations in the delta frequency range are shown. These associations between Post-learning channel-pair similarity in the delta frequency and statistical consolidation were more pronounced in more adjacent channel-pairs. On the left of each panel, the mean absolute values of correlation coefficients for the short (lowest 25% IQR), medium (middle 25% IQR), and long (highest 25% IQR) distance of the electrodes in the channel-pairs (Euclidean distance of the Cartesian coordinates of the electrodes) for serial-order (A) and statistical (B) consolidation are depicted. The error bars indicate the SEM. On the right of each panel, linear trendlines for the distance of the electrodes in the channel-pairs and the correlation strength of the similarities of those channel-pairs with the consolidation index of serial-order (A) and statistical (B) knowledge are shown, separately for positive (lighter color) and negative (darker color) correlations. The dots indicate individual correlation coefficients for each channel-pair. \*\*\**p* < .005, ** *p* < .01, +*p* < .1

The ANOVA on the correlation coefficients between the consolidation index of statistical knowledge and the channel-pair similarity values in the delta frequency range as a function of the distance between the electrodes of the channel-pairs was also significant: (F_2,761_ = 5.49, *p* = .004, η_p_^2^ = .009): absolute values of correlation coefficients were different as a function of distance (Fig. 8B, left). The pairwise post-hoc comparisons revealed that channel-pairs with long distances showed significantly weaker, and a trend for weaker correlations with statistical consolidation than channel-pairs with short and medium distances respectively (*p =* .004, *p* = .09). Correlation strengths did not differ between short and medium distances (*p* = .26). When considering the direction of correlations (positive or negative correlation coefficients), we found that the trend visible in the ANOVA, namely that statistical consolidation showed stronger associations with shorter connections, was led by the negative correlation coefficients (Fig. 8B, right).

#### Trait-like associations between behavior and similarity in EEG functional connectivity across different periods

To disentangle learning-induced vs. trait-like similarities, we compared Baseline and Postl-earning similarity for frequency bins and their associations with the consolidation indices. If the Baseline similarity (that is, the similarity of functional connectivity during learning and pre-learning rest) and its relation to memory consolidation is comparable to Post-learning similarity (that is, the similarity of functional connectivity during learning and post-learning rest), the resemblance may have emerged from stable individual characteristics in brain activity. Figure 5A shows similar patterns for Baseline and Post-learning similarity computed for frequency bins. To compare the associations of Baseline and Post-learning similarity, we conducted the same correlation analyses with the consolidation indices for the raw similarity values as we computed for the Learning-induced similarity change values in the ‘Frequency of reactivation’ section of the Results. The consolidation of serial-order knowledge showed a significant positive correlation in the alpha frequency range (for the 9.5-10.5 Hz bins) with Post-learning similarity (Fig. 9A). The pattern of the associations of the consolidation of serial-order knowledge was similar over the Baseline and the Post-learning similarity measures, however, the effects were weaker and not significant for the Baseline similarity in the alpha frequency range (Fig. 9A). The consolidation of statistical knowledge showed a significant negative correlation in the delta frequency range (2.5-3 Hz) with the Baseline similarity (Fig. 9B). Again, Baseline and Post-learning similarities exhibited similar patterns, however, the effects were weaker and not significant for the Post-learning similarity in the delta frequency range (Fig. 9B). To sum up, the similar pattern of the Baseline and Post-learning similarity per se (Fig. 5) and the similar patterns of their associations with the consolidation indices (Fig. 9) suggest that trait-like associations exist between the brain activity during different periods and memory consolidation.

**Figure 9.**
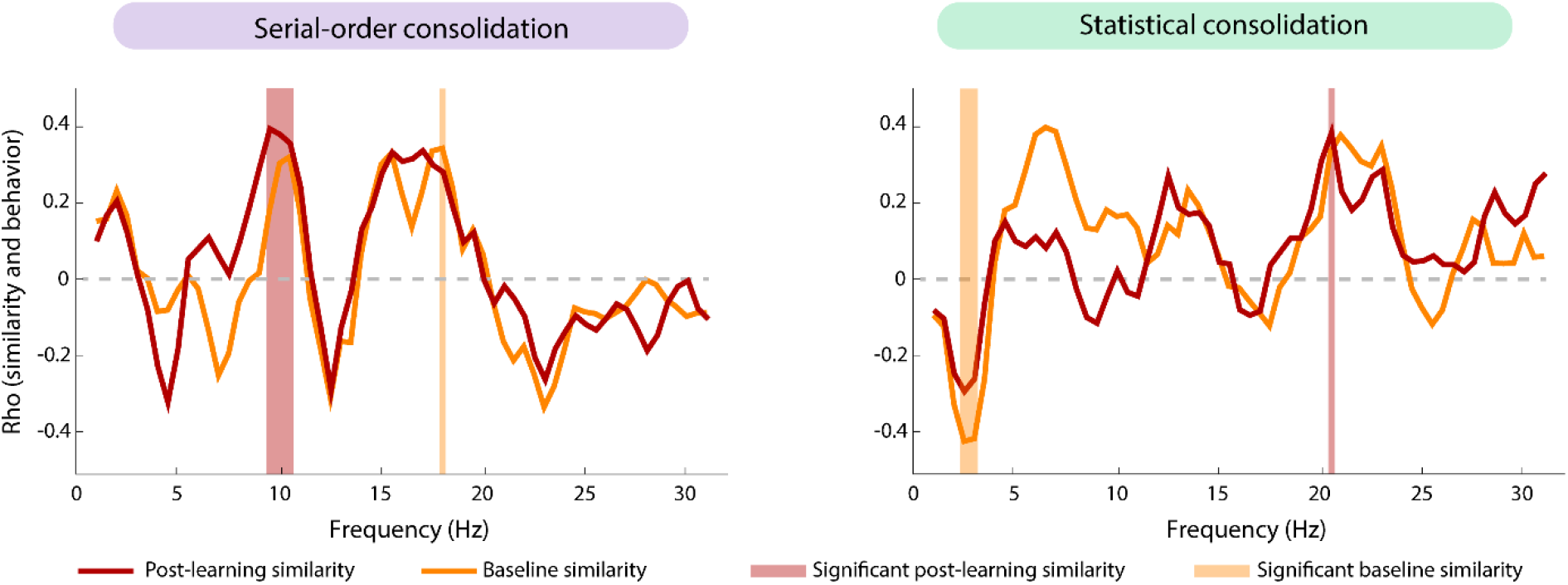
Correlation coefficients (Spearman Rho’s) of Baseline (orange) and Post-learning (red) similarity and the consolidation indices of serial-order (left) and statistical (right) knowledge. The patterns of the associations of the consolidation of serial-order and statistical knowledge were similar over the Baseline and the Post-learning similarity measures. The shading denotes the significant (*p* < .05) correlations after cluster-based correction for multiple comparisons.

Besides the trait-like associations, these analyses also unraveled the dynamics of the associations of the Learning-induced similarity changes in more detail: The positive correlation between the Learning-induced similarity change values and memory consolidation (Fig. 6) occurred for different reasons in alpha and delta frequency. In the alpha frequency, both the Baseline and the Post-learning similarity values showed positive correlations with the consolidation of serial-order knowledge, however, it was stronger for the Post-learning similarity. In contrast, in the delta frequency, both the Baseline and the Post-learning similarity values showed negative correlations with the consolidation of statistical knowledge, however, it was weaker for the Post-learning similarity.

#### Controlling for the influence of the raw functional connectivity

Importantly, the associations between the consolidation indices and similarity/learning-induced similarity change values are not caused by the functional connectivity of the learning (or pre- or post-learning rest) period, as the correlation between the raw connectivity matrices during learning (or pre- or post-learning rest) and serial-order and statistical consolidation showed different patterns (Fig. 10).

**Figure 10.**
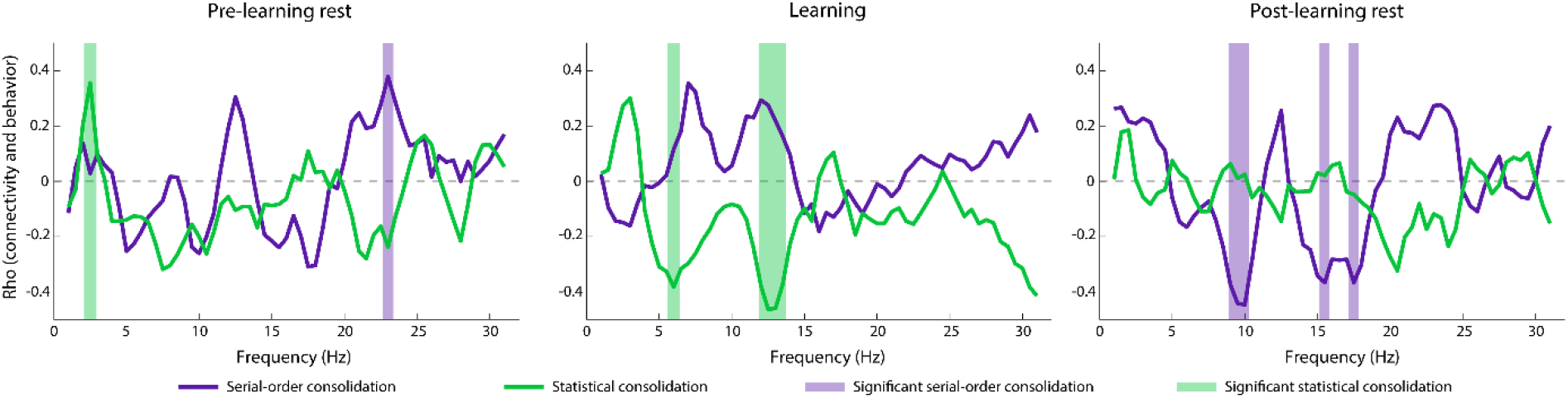
Correlation coefficients (Spearman Rho’s) between behavioral indices of consolidation and EEG functional connectivity measured as the Weighted Phase Lag Index (WPLI) during pre-learning resting state (left), learning (middle) and post-learning resting state (right). Purple and green lines indicate associations with the consolidation of serial-order and statistical knowledge, respectively. The shading denotes the significant (*p* < .05) correlations after cluster-based correction for multiple comparisons.

## Discussion

Our results showed that higher similarity in functional connectivity patterns between learning and post-learning rest compared to pre-learning rest (termed Learning-induced similarity change) in the alpha frequency range (8.5-9.5 Hz) is beneficial for the consolidation of serial-order knowledge. At the same time, Learning-induced similarity change in the delta frequency range (2.5-3 Hz) was positively associated with the consolidation of statistical (probability-based) knowledge. The topographical analyses within these frequency ranges highlighted the involvement of long-range centro-parietal connections in the consolidation of serial-order knowledge, and shorter-range connections in the consolidation of statistical knowledge.

Beyond these learning-induced changes, we also compared the raw similarity matrices and their associations with memory consolidation to explore trait-like similarities. Similarities in functional connectivity of learning and post-learning rest (Post-learning similarity) and learning and pre-learning rest (Baseline similarity) exhibited similar patterns, and showed comparable associations with the consolidation indices, indicating the existence of trait-like resemblance. Furthermore, these comparisons shed light on two different dynamics underlying the positive correlations between the Learning-induced similarity change values and the consolidation indices: in case of serial-order knowledge, better consolidation was associated with the strengthening of a positive correlation (beneficial effect) between the similarity values and the consolidation index from Baseline to Post-learning similarity, whereas in case of statistical knowledge, better consolidation was associated with the weakening of a negative correlation (detrimental effect).

Our results suggest that learning-induced changes in the alpha synchronization are beneficial for the consolidation of serial-order (but not probability) information (Fig. 6). How did alpha synchronization change over the different periods and what part of these changes were relevant for memory consolidation? Concerning the former, alpha synchronization was higher during both resting states than during learning and was higher during post-learning rest than during the pre-learning rest (Fig. 4). Similarly to our study, Murphy et al. (2018) showed higher synchrony in alpha frequency during post-learning rest compared to pre-learning rest. Importantly, this was only true when there was a learning episode involved between the two resting states (not when other cognitive tasks were completed). Concerning the relevant changes in alpha synchronization, the beneficial (higher) similarity between learning and post-learning rest could occur due to higher alpha synchronization during the learning or lower alpha synchronization during post-learning rest. Based on the correlation of the raw functional connectivity of learning and post-learning rest with memory consolidation showing that connectivity in the alpha frequency of post-learning rest negatively correlates with the consolidation of serial-order knowledge (Fig. 10), the associations with the similarity (Fig. 6) are most likely led by lower alpha synchronization during the post-learning rest. This is in line with the study of Brokaw et al. (2016), which showed that greater alpha power during rest following a declarative learning task was negatively correlated with subsequent recall performance. In other words, higher alpha synchronization was detrimental during the post-learning rest period for memory retention. Altogether, these results suggest that alpha synchronization increases after learning, but individuals showing smaller increase in alpha synchronization have better memory consolidation.

Importantly, similarity in functional networks can occur not only due to generally higher or lower synchronization but dynamics in synchronization that are present during different periods. The functional role of the alpha rhythm (i.e., excitatory (Palva and Palva, 2007) or inhibitory (Klimesch et al., 2007; Jensen and Mazaheri, 2010)) seems to depend on phase dynamics (Palva and Palva, 2011), firing modes (Peterson and Voytek, 2017) and behavioral states (Alamia and VanRullen, 2019). Thus, similarity in alpha synchronization during learning and subsequent resting-state could reflect reactivated or maintained brain dynamics in some of these factors.

Why learning-induced changes beneficial for the consolidation of serial-order information were found in alpha frequency? Von Stein and Sarnthein (2000) suggested that the longer the range of the cortical connections, the slower the oscillatory frequency is through which they synchronize; that is, alpha as a relatively slow oscillation has been proposed to have a large-scale integrative function. Based on the tasks that involve higher alpha synchronization (e.g., mental imagery, working memory, resting state), they suggested that greater alpha synchronization reflects internal mental activity and may be responsible for top-down cognitive processes. Top-down here refers to the flow of information processing: for top-down processes, the sensory processing is influenced by higher-order cognition, i.e., it is model-dependent/hypothesis-driven, while for bottom-up processes, sensory processing is driven by the stimuli, i.e., it is model-free. In support of the role of alpha in top-down processing, a recent paper showed that alpha propagates from higher- to lower-order cortex and cortex to thalamus (Halgren et al., 2019), providing a laminar circuit for top-down processes. Our results are in line with the findings that show the role of alpha in top-down processes as we demonstrated that synchronization in alpha frequency is important for serial-order memory consolidation, which in our study was a top-down, attention-demanding process, while it did not show association with statistical learning, which is a bottom-up, stimulus-driven process (Nemeth et al., 2013). The topographical distribution of relevant phase-synchronization in alpha frequency (Fig. 7-8) is also consistent with the integrative/top-down function of alpha oscillation in serial-order memory consolidation: Based on the channel-pair similarity correlations in alpha frequency, our results indicate that long-range centro-parietal connections are important for this type of memory. This is also in line with previous studies investigating the neural background of serial-order learning that seems to at least partially rely on fronto-parietal networks (Sakai et al., 1998; Doyon et al., 2009).

In contrast to the consolidation of serial-order knowledge, the consolidation of statistical knowledge showed associations with the learning-induced changes in the delta frequency range. However, this change was a weakening detrimental effect, i.e. decreasing adverse association between the consolidation of statistical knowledge and similarity from Baseline to Post-learning similarity. Why learning-induced changes beneficial for the consolidation of probability information were found in delta frequency? Delta oscillation during cognitive tasks has been proposed to play a role in inhibiting interferences that may affect task performance (Harmony, 2013). Together with the topography of connections that correlated (negatively) with statistical learning, this suggests that the inhibition of the left, in particular parietal, sites is beneficial for statistical memory. This is in line with studies showing that the right hemisphere is important for statistical learning (Roser et al., 2011; Shaqiri and Anderson, 2013; Janacsek et al., 2015).

The learning-induced changes assessed in our study could emerge partly due to reactivation. However, the definition of reactivation should be understood broadly in our study, as the similarities discussed here could occur as a result of reactivation of the memory traces per se, the reactivation of the overall learning experience, or the reactivation of the cognitive set in general during the learning episode.

Lastly, our results draw attention to the great overlap of active functional networks during a task and preceding/subsequent resting states. These similarities during different behavioral states have been largely neglected in previous studies investigating reactivation (as reactivation is traditionally defined by the unique overlap between the learning and the post-learning off-line period). However, the functional brain networks that are present during other behavioral states might also be important for memory consolidation. We argue that not only reoccurring but also stable, trait-like brain networks are important for cognitive performance, particularly for memory consolidation. For example, studies have shown that there is little difference between functional connectivity networks during resting state and a task measured via fMRI (Cole et al., 2016; Satterthwaite et al., 2018). Gratton et al. (2018) concluded that functional networks are dominated by stable individual factors, not cognitive content. Our results are in line with these fMRI results and demonstrate that these trait-like, stable networks can be captured by EEG as well and they support memory consolidation.

### Conclusions

Our results show that learning-induced changes in resting functional networks after learning differentially predict memory consolidation of serial-order and probability information acquired simultaneously: learning-induced changes in alpha synchronization were associated with the consolidation of serial-order, whereas learning-induced changes in delta synchronization were associated with the consolidation of probability information. Long-range functional connections in the alpha frequency seem to be important for the consolidation of serial-order knowledge, potentially due to the involvement of top-down processes in this type of learning, requiring the synchronization of distant brain areas. In contrast, consolidation of statistical (probability-based) knowledge seems to rely on local synchronization, potentially due to the involvement of bottom-up, stimulus-driven processes that are less dependent on long-range synchronization. Finally, patterns of similarities between learning and pre- and post-learning rest (and their associations with consolidation performance) showed great overlap. This implies that rather than, or in parallel with, reoccurring brain dynamics, maintained networks are also important for cognitive performance, particularly for memory consolidation. Lastly, we provided a framework for studying memory consolidation in its complexity, incorporating the simultaneous consolidation of different information and the influence of learning-induced changes and stable characteristics of brain activity on the consolidation performance.

## Acknowledgements

This research was supported by the Research and Technology Innovation Fund, Hungarian Brain Research Program (National Brain Research Program, project 2017-1.2.1-NKP-2017-00002, to DN); the IDEXLYON Fellowship of the University of Lyon as part of the Programme Investissements d’Avenir (ANR-16-IDEX-0005, to DN); the Hungarian Scientific Research Fund (NKFIH-OTKA PD 124148, to KJ; NKFIH-OTKA K 128016, to DN); the National Research, Development and Innovation Office (NKFIH FK 128100, NKFIH-1157-8/2019-DT, to PS) and Janos Bolyai Research Fellowship of the Hungarian Academy of Sciences (to KJ). We thank Rafael Pedrosa for his invaluable help with the graphical illustrations.

## References

Alamia A, VanRullen R (2019) Alpha oscillations and traveling waves: Signatures of predictive coding? PLoS Biol 17.

Brokaw K, Tishler W, Manceor S, Hamilton K, Gaulden A, Parr E, Wamsley EJ (2016) Resting state EEG correlates of memory consolidation. Neurobiol Learn Mem 130:17–25.

Buzsáki G, Draguhn A (2004) Neuronal oscillations in cortical networks. Science 304:1926–1929.

Cohen MX (2014) Analyzing neural time series data: theory and practice: MIT press.

Cole MW, Ito T, Bassett DS, Schultz DH (2016) Activity flow over resting-state networks shapes cognitive task activations. Nat Neurosci 19:1718.

Doyon J, Bellec P, Amsel R, Penhune V, Monchi O, Carrier J, Lehericy S, Benali H (2009) Contributions of the basal ganglia and functionally related brain structures to motor learning. Behavioral Brain Research 199:61–75.

Gratton C, Laumann TO, Nielsen AN, Greene DJ, Gordon EM, Gilmore AW, Nelson SM, Coalson RS, Snyder AZ, Schlaggar BL (2018) Functional brain networks are dominated by stable group and individual factors, not cognitive or daily variation. Neuron 98:439–452. e435.

Greenhouse SW, Geisser S (1959) On methods in the analysis of profile data. Psychometrika 24:95–112.

Halgren M, Ulbert I, Bastuji H, Fabó D, Erőss L, Rey M, Devinsky O, Doyle WK, Mak-McCully R, Halgren E (2019) The generation and propagation of the human alpha rhythm. Proceedings of the National Academy of Sciences 116:23772–23782.

Harmony T (2013) The functional significance of delta oscillations in cognitive processing. Front Integr Neurosci 7:83.

Hermans EJ, Kanen JW, Tambini A, Fernández G, Davachi L, Phelps EA (2017) Persistence of amygdala–hippocampal connectivity and multi-voxel correlation structures during awake rest after fear learning predicts long-term expression of fear. Cereb Cortex 27:3028–3041.

Howard JH, Jr., Howard DV (1997) Age differences in implicit learning of higher-order dependencies in serial patterns. Psychol Aging 12:634–656.

Janacsek K, Ambrus GG, Paulus W, Antal A, Nemeth D (2015) Right hemisphere advantage in statistical learning: evidence from a probabilistic sequence learning task. Brain stimulation 8:277–282.

Jensen O, Mazaheri A (2010) Shaping functional architecture by oscillatory alpha activity: gating by inhibition. Front Hum Neurosci 4:186.

Klimesch W, Sauseng P, Hanslmayr S (2007) EEG alpha oscillations: the inhibition–timing hypothesis. Brain research reviews 53:63–88.

Lee T-W, Girolami M, Sejnowski TJ (1999) Independent component analysis using an extended infomax algorithm for mixed subgaussian and supergaussian sources. Neural Comput 11:417–441.

Louie K, Wilson MA (2001) Temporally structured replay of awake hippocampal ensemble activity during rapid eye movement sleep. Neuron 29:145–156.

Maheu M, Meyniel F, Dehaene S (2020) Rational arbitration between statistics and rules in human sequence learning. bioRxiv.

Maquet P, Laureys S, Peigneux P, Fuchs S, Petiau C, Phillips C, Aerts J, Del Fiore G, Degueldre C, Meulemans T, Luxen A, Franck G, Van Der Linden M, Smith C, Cleeremans A (2000) Experience-dependent changes in cerebral activation during human REM sleep. Nat Neurosci 3:831–836.

McGaugh JL (2000) Memory--A century of consolidation. Science 287:248–251.

McGaugh JL, Izquierdo I (2000) The contribution of pharmacology to research on the mechanisms of memory formation. Trends Pharmacol Sci 21:208–210.

Murphy M, Stickgold R, Parr ME, Callahan C, Wamsley EJ (2018) Recurrence of task-related electroencephalographic activity during post-training quiet rest and sleep. Sci Rep 8:1–10.

Nádasdy Z, Hirase H, Czurkó A, Csicsvari J, Buzsáki G (1999) Replay and time compression of recurring spike sequences in the hippocampus. J Neurosci 19:9497–9507.

Nemeth D, Janacsek K, Fiser J (2013) Age-dependent and coordinated shift in performance between implicit and explicit skill learning. Front Comput Neurosci 7.

Palva S, Palva JM (2007) New vistas for α-frequency band oscillations. Trends Neurosci 30:150–158.

Palva S, Palva JM (2011) Functional roles of alpha-band phase synchronization in local and large-scale cortical networks. Front Psychol 2:204.

Peigneux P, Orban P, Balteau E, Degueldre C, Luxen A, Laureys S, Maquet P (2006) Offline persistence of memory-related cerebral activity during active wakefulness. PLoS Biol 4:e100.

Peigneux P, Laureys S, Fuchs S, Collette F, Perrin F, Reggers J, Phillips C, Degueldre C, Del Fiore G, Aerts J (2004) Are spatial memories strengthened in the human hippocampus during slow wave sleep? Neuron 44:535–545.

Peterson EJ, Voytek B (2017) Alpha oscillations control cortical gain by modulating excitatory-inhibitory background activity. Biorxiv:185074.

Peyrache A, Khamassi M, Benchenane K, Wiener SI, Battaglia FP (2009) Replay of rule-learning related neural patterns in the prefrontal cortex during sleep. Nat Neurosci 12:919.

Qin Y-L, McNaughton BL, Skaggs WE, Barnes CA (1997) Memory reprocessing in corticocortical and hippocampocortical neuronal ensembles. Philos Trans R Soc Lond B Biol Sci 352:1525–1533.

Rasch B, Born J (2007) Maintaining memories by reactivation. Curr Opin Neurobiol 17:698–703.

Ribeiro S, Gervasoni D, Soares ES, Zhou Y, Lin S-C, Pantoja J, Lavine M, Nicolelis MA (2004) Long-lasting novelty-induced neuronal reverberation during slow-wave sleep in multiple forebrain areas. PLoS Biol 2.

Rickard TC, Cai DJ, Rieth CA, Jones J, Ard MC (2008) Sleep does not enhance motor sequence learning. Journal of Experimental Psychology: Learning, Memory, and Cognition; Journal of Experimental Psychology: Learning, Memory, and Cognition 34:834.

Rieth CA, Cai DJ, McDevitt EA, Mednick SC (2010) The role of sleep and practice in implicit and explicit motor learning. Behav Brain Res 214:470–474.

Roser ME, Fiser J, Aslin RN, Gazzaniga MS (2011) Right hemisphere dominance in visual statistical learning. J Cogn Neurosci 23:1088–1099.

Rothschild G, Eban E, Frank LM (2017) A cortical–hippocampal–cortical loop of information processing during memory consolidation. Nat Neurosci 20:251–259.

Sakai K, Hikosaka O, Miyauchi S, Takino R, Sasaki Y, Putz B (1998) Transition of brain activation from frontal to parietal areas in visuomotor sequence learning. J Neurosci 18:1827–1840.

Satterthwaite TD, Xia CH, Bassett DS (2018) Personalized neuroscience: Common and individual-specific features in functional brain networks. Neuron 98:243–245.

Shaqiri A, Anderson B (2013) Priming and statistical learning in right brain damaged patients. Neuropsychologia 51:2526–2533.

Simor P, Zavecz Z, Horvath K, Elteto N, Török C, Pesthy O, Gombos F, Janacsek K, Nemeth D (2019) Deconstructing procedural memory: Different learning trajectories and consolidation of sequence and statistical learning. Front Psychol 9:2708.

Skaggs WE, McNaughton BL (1996) Replay of neuronal firing sequences in rat hippocampus during sleep following spatial experience. Science 271:1870.

Song S, Howard JH, Jr., Howard DV (2007) Implicit probabilistic sequence learning is independent of explicit awareness. Learn Mem 14:167–176.

Stam CJ, Nolte G, Daffertshofer A (2007) Phase lag index: assessment of functional connectivity from multi channel EEG and MEG with diminished bias from common sources. Hum Brain Mapp 28:1178–1193.

Tambini A, Davachi L (2019) Awake Reactivation of Prior Experiences Consolidates Memories and Biases Cognition. Trends in cognitive sciences.

Tambini A, Berners-Lee A, Davachi L (2017) Brief targeted memory reactivation during the awake state enhances memory stability and benefits the weakest memories. Sci Rep 7:1–17.

Varela F, Lachaux J-P, Rodriguez E, Martinerie J (2001) The brainweb: phase synchronization and large-scale integration. Nat Rev Neurosci 2:229–239.

Vinck M, Oostenveld R, Van Wingerden M, Battaglia F, Pennartz CM (2011) An improved index of phase-synchronization for electrophysiological data in the presence of volume-conduction, noise and sample-size bias. Neuroimage 55:1548–1565.

Von Stein A, Sarnthein J (2000) Different frequencies for different scales of cortical integration: from local gamma to long range alpha/theta synchronization. Int J Psychophysiol 38:301–313

